# Prim-O-glucosylcimifugin suppresses M2 macrophage polarization via the JAK2-STAT6-PPARγ pathway to alleviate allergic asthma

**DOI:** 10.64898/2026.11.10.765782

**Authors:** Chong Li, Hong Zhang, Weiwei Yang, Haiyan Fu, Honghong Huang

## Abstract

**Background:** Allergic asthma (AA) is a chronic respiratory disease closely linked to immune-inflammatory responses driven by M2 macrophage polarization. PrimO-glucosylcimifugin (POG), a natural bioactive compound from Saposhnikovia divaricata (SD), alleviates inflammation in acute lung injury mice, but whether it mitigates AA by modulating M2 macrophage polarization remains unclear.

**Objective:** This study investigated POG’s effects on AA and its mechanisms in regulating M2 macrophage polarization.

**Methods:** LC-MS profiled the SD/Prunus mume (PM) combination, and network pharmacology predicted its main active components and key anti-AA mechanisms. Ovalbumin (OVA) plus IL-4 established AA models in mice and RAW264.7 cells. Lung protection was assessed by histopathology, lung indices, and type II immune inflammatory cytokine markers. Flow Cytometry, RT-qPCR, immunofluorescence, and Western blot examined key genes/proteins in the JAK2-STAT6-PPARγ axis.

**Results:** LC-MS-based network pharmacology identified POG as the key SD/PM active ingredient alleviating AA. Pharmacodynamic evaluation showed POG effectively alleviated OVA-induced airway inflammation and pulmonary histopathological damage in AA mice. Mechanistically, POG inhibited M2 macrophage polarization by downregulating key JAK2-STAT6-PPARγ axis factors.

**Conclusion:** POG ameliorates AA by reducing type II immune factor release through JAK2-STAT6-PPARγ-mediated inhibition of M2 macrophage polarization. This study clarifies POG’s anti-AA mechanism and supports its clinical development.

## 1 Introduction

Allergic asthma (AA) is the most common subtype of bronchial asthma, characterized by chronic airway inflammation and airway hyperresponsiveness [1]. Patients often present with recurrent symptoms such as wheezing, shortness of breath, chest tightness, or coughing [2]. Approximately 300 million people worldwide suffer from asthma, and the prevalence of AA is rising year by year, placing an increasingly heavy economic burden on society [3]. Currently, clinical treatment strategies primarily involve the use of corticosteroids [4]. However, significant adverse drug reactions and a high risk of developing drug resistance during treatment severely limit clinical efficacy[5]. Therefore, there is an urgent need to identify a safer and more effective treatment strategy, which holds significant practical importance for both the clinical management of AA and the alleviation of the societal burden.

In the onset and progression of AA, macrophage polarization plays a key role in the body’s immune and inflammatory responses [6]. This polarization process is finely regulated by signals from the local microenvironment [7]. As a key component of the lung’s innate defense mechanism, macrophages can polarize into either the M1 or M2 immune phenotypes [8]. Studies have shown that M2 macrophage polarization promotes the release of type II cytokines such as IL-5, IL-4, and IL-13, thereby amplifying the pulmonary inflammatory response and exacerbating airway hyperresponsiveness [9]. Another study confirmed that the JAK2-STAT6-PPARγ signaling axis is a key upstream pathway driving M2 phenotypic polarization [10]. In this signaling axis, upstream-activated JAK2 phosphorylates STAT6; the activated STAT6 then directly binds to and activates the DNA promoter of the PPAR-γ gene, promoting its transcriptional expression [10]. PPAR-γ is a nuclear receptor transcription factor, and its transcriptional activity is a key factor in the polarization of macrophages toward the M2 phenotype [11]. Therefore, inhibiting M2 macrophage polarization mediated by the JAK2-STAT6-PPAR-γ axis, thereby reducing the excessive release of cytokines, may represent an important strategy for alleviating AA.

Traditional Chinese medicine (TCM) offers the advantage of multi-target, holistic regulation in the treatment of asthma, with its anti-inflammatory and immunomodulatory effects drawing particular attention [12,13]. The combination of *Saposhnikovia divaricata* (SD)/*Prunus mume* (PM), derived from the “Allergy Decoction” formulated by the renowned traditional Chinese medicine physician Zhanyu Zhu, possesses pharmacological effects such as anti-inflammatory, antioxidant, and immunomodulatory actions, and is commonly used to treat bronchial asthma [14]. Modern research indicates that SD/PM exerts anti-allergic effects by blocking mast cell degranulation and downregulating the Akt/ERK signaling pathway [15]. Our previous studies have also confirmed that SD/PM can alleviate airway inflammation in AA mice by regulating the Th1/Th2 immune balance [16]. Building on this, the present study screened and identified PrimO-glucosylcimifugin (POG) as the key active component in SD/PM responsible for its therapeutic effects on AA through the integration of LC-MS identification and network pharmacology analysis. POG is one of the most abundant chromone compounds in FF. Previous studies have demonstrated that it can alleviate liver injury by inhibiting NLRP3 inflammasome-mediated macrophage activation and exert a protective effect in LPS-induced acute lung injury by regulating the MAPK/NF-κB pathway[14, 15]. However, the role of POG in AA remains unclear, and whether it exerts anti-AA effects by modulating M2 macrophage polarization through the JAK2-STAT6-PPARγ axis still needs to be elucidated.

This study focused on POG and systematically investigated its therapeutic effects on AA and the underlying molecular mechanisms by establishing an Ovalbumin (OVA)-induced AA mouse model and an IL-4-induced RAW264.7 macrophage polarization model. In this study, we demonstrated that POG effectively alleviates airway inflammation and pulmonary pathological damage in AA mice. Mechanistically, POG exerts its beneficial effects on AA by inhibiting M2 macrophage polarization mediated by the JAK2-STAT6-PPARγ axis. This study indicates that POG may serve as a candidate drug for the treatment of AA, which is of great significance for curbing the progression of the disease and alleviating the socioeconomic and healthcare burdens.

## 2 Materials and methods

### 2.1 Experimental drugs and main reagents

The herbal materials SD and PM were both purchased from Haikou Traditional Chinese Medicine Hospital. OVA (Batch No. A5503) was purchased from Sigma-Aldrich. The hematoxylin-eosin staining kit (Batch No. G1005) and the PAS staining kit (Batch No. G1008) were purchased from Servicebio Co., Ltd. The Masson staining kit (Batch No. G1340) was purchased from Solarbio Co., Ltd. The IgE (Batch No. PI476), IL-4 (Batch No. PI612), IL-5 (Batch No. PI620), ELISA kits for IL-13 (Batch No. PI539) were purchased from Biyuntian Biological Co., Ltd. Trizol, Evo M-MLV RT Mix Kit with gDNA Clean for qPCR, and SYBR® Green Premix Pro Taq HS qPCR Kit were purchased from AG Scientific Co., Ltd. The Senescence *β*-Galactosidase Staining Kit (Batch No. C0602) was purchased from Biyuntian Biological Co., Ltd. CD86 (Batch No. BF3037), CD206 (Batch No. BF3017), and F4/80 (Batch No. BF3097) were purchased from Affinity Biosciences. The JAK2 antibody (Batch No. TA6022), p-JAK2 antibody (Batch No. PC3206), STAT6 antibody (Batch No. T55351), p-STAT6 antibody (Batch No. TB3941), PPARγ antibody (Batch No. PC0530), and p-PPARγ antibody (Batch No. PC3737) were purchased from Abmart.

### 2.2 Preparation of SD/PM Extracts

Weigh out 15 g of SD and PM, add 10 times the weight (w/v) in ultrapure water, soak for 30 minutes, then heat and boil for 20 minutes, and collect the filtrate. Add 8 times the volume of water to the herbal residue and continue boiling twice, for 20 minutes each time; combine the filtrates. Combine the resulting filtrates and concentrate them under reduced pressure to a thick paste; then subject them to vacuum freeze-drying to obtain the freeze-dried powder. Store the freeze-dried powder at 4°C for future use; reconstitute with ultrapure water immediately before use.

### 2.3 Animal experiments

This study was approved by the Medical Ethics Committee of Haikou Traditional Chinese Medicine Hospital (Review Approval No.: HHTCM2025-01-08). Thirty-five 8-week-old SPF-grade male BALB/c mice were randomly divided into 7 groups (n = 5): control group, model group, dexamethasone group (1 mg/kg, positive control), low-dose POG group (12.5 mg/kg), the medium-dose POG group (25 mg/kg), the high-dose POG group (50 mg/kg), and the high-dose POG group + Coumermycin A1 (C-A1, 1 mg/kg). On days 1, 7, and 14, mice in all groups except the control group were sensitized via intraperitoneal injection with an emulsion of ovalbumin (OVA, 100 μg) and aluminum hydroxide adjuvant (2 mg); the control group received an equal volume of saline. On days 15–21, the dexamethasone group received daily intraperitoneal injections of dexamethasone (1 mg/kg), while the POG groups at each dose received daily intraperitoneal injections of the corresponding concentration of POG solution. The control and model groups received an equal volume of saline for 7 consecutive days. On the 22nd day of the experiment, all mice were anesthetized and euthanized; blood and lung tissue were collected and stored at -80°C for future use.

### 2.4 LC-MS Analysis

The SD/PM extract was centrifuged at 12,000 rpm for 15 min at 4°C. The supernatant was collected and analyzed using a Thermo Fisher Scientific Vanquish ultra-high-performance liquid chromatography system with a C18 column (1.7 μm, 2.1 × 100 mm, Waters). The mobile phase flow rate was maintained at 0.35 mL/min, and the injection volume was 5 μL. The mobile phase consisted of an aqueous phase (A) containing 0.1% formic acid and an acetonitrile phase (B) containing 0.5% formic acid. Compound identification was confirmed by comparison with the in-house BiotreeDB secondary mass spectrometry library (V2.1).

### 2.5 Network Pharmacology Analysis

Compounds contained in “*Saposhnikovia divaricata*” and “*Prunus mume*” were retrieved from the TCMSP database (https://old.tcmsp-e.com/tcmspsearch.Php) and screened based on the criteria of oral bioavailability (OB) ≥ 20% and drug-likeness (DL) ≥ 0.18[17]. A Venn diagram was created by overlaying the screened compounds with those identified by LC-MS; the intersection represents the candidate compounds for SD/PM. The potential drug targets for these candidate compounds were predicted using SwissTargetPrediction (http://www.swisstargetprediction.ch/). Simultaneously, “Allergic Asthma” was searched in the GeneGards (https://auth.lifemapsc.com/) and OMIM (https://omim.org/) databases to identify candidate targets. Map the targets related to “Allergic Asthma” to the drug targets and create a Venn diagram to display the overlapping targets. Import the overlapping targets into the STRING database (https://cn.string-db.org/) for PPI analysis, then visualize the results using Cytoscape 3.10.1. Use the DAVID database (https://davidbioinformatics.nih.gov) to perform GO and KEGG analyses on the overlapping targets, and create a bubble chart displaying functional and pathway information. Based on this, construct a “drug–component– pathway–target–disease” network to illustrate the interactions among drug components, targets, and diseases.

### 2.6 Pathological Evaluation of Lung Tissue

Lung tissue was collected from mice in each group, preserved in a fixative containing 4% paraformaldehyde, dehydrated with ethanol, embedded, and sectioned into 4-μm-thick sections. The sections were then stained with hematoxylin and eosin (H&E), PAS, and Masson’s stain, scanned using a digital microtome scanner, and the pathological features of the mouse lung tissue were observed and imaged.

### 2.7 ELISA

Blood samples were collected from the mice and allowed to clot at room temperature, followed by centrifugation at 3,000 × g for 10 min at 4 °C. The serum was carefully collected and stored at −80 °C until further analysis. The levels of total IgE, IL-4, IL-5, and IL-13 in the serum were determined using commercially available ELISA kits according to the manufacturer’s instructions. The absorbance values were measured using a microplate reader, and the concentrations of the target proteins were calculated according to the corresponding standard curves.

### 2.8 Cell Culture

Raw264.7 cells were obtained from Haixing Biotechnology Co., Ltd. The cells were cultured in DMEM medium containing 10% FBS, 1% penicillin, and 1% streptomycin at 37 °C in a 5% CO_2_ atmosphere. Raw264.7 cells were treated with 25 ng/mL IL-4 for 24 hours to induce M2 polarization. When the cells reached 80% confluence, they were passaged at a 1:2 ratio.

### 2.9 Cell Viability Assay

RAW264.7 cells were seeded at an appropriate density per well in a 96-well plate and allowed to adhere for 24 hours in an incubator at 37 °C and 5% CO_2_. After stimulating the cells with 25 ng/mL IL-4 for 24 hours, the supernatant was removed, and fresh complete medium containing gradient concentrations of POG was added, and the cells were cultured for 24 hours, respectively. CCK-8 reagent was then added to each well; after incubation, OD values were measured at 450 nm, and cell viability was expressed as a percentage relative to the control group.

### 2.10 Flow cytometry

After grinding the lung tissue and centrifuging it, the supernatant was discarded, and the cell pellet was collected and set aside. RAW264.7 cells were treated as appropriate and then washed twice with staining buffer. Subsequently, the cells were incubated with fluorescently labeled anti-CD86 and anti-CD206 antibodies for 40 minutes at 4 °C in the dark. After staining, the cells were washed twice with PBS, resuspended in an appropriate volume of staining buffer, and immediately analyzed by flow cytometry. Unstained controls, single-stain controls, and isotype controls were included in the experiment to establish gating strategies and adjust fluorescence compensation.

### 2.11 Reverse-transcription quantitative polymerase chain reaction (RT-qPCR)

Total RNA was extracted from mouse lung tissues and cells using Trizol reagent, and the RNA concentration was determined using a nucleic acid analyzer. Subsequently, the RNA was reverse-transcribed into cDNA using the Evo M-MLV RT Mix Kit with gDNA Clean for qPCR under the following conditions: 95 °C for 30 s, followed by 40 cycles of 95 °C for 10 s and 60 °C for 30 s. The relative expression levels of CD206, Arg-1, and GAPDH mRNAs in tissues and cells were determined using the SYBR Premix Ex Taq™ II Kit. The primer sequences are shown in **Table 1**.

**Table 1.**
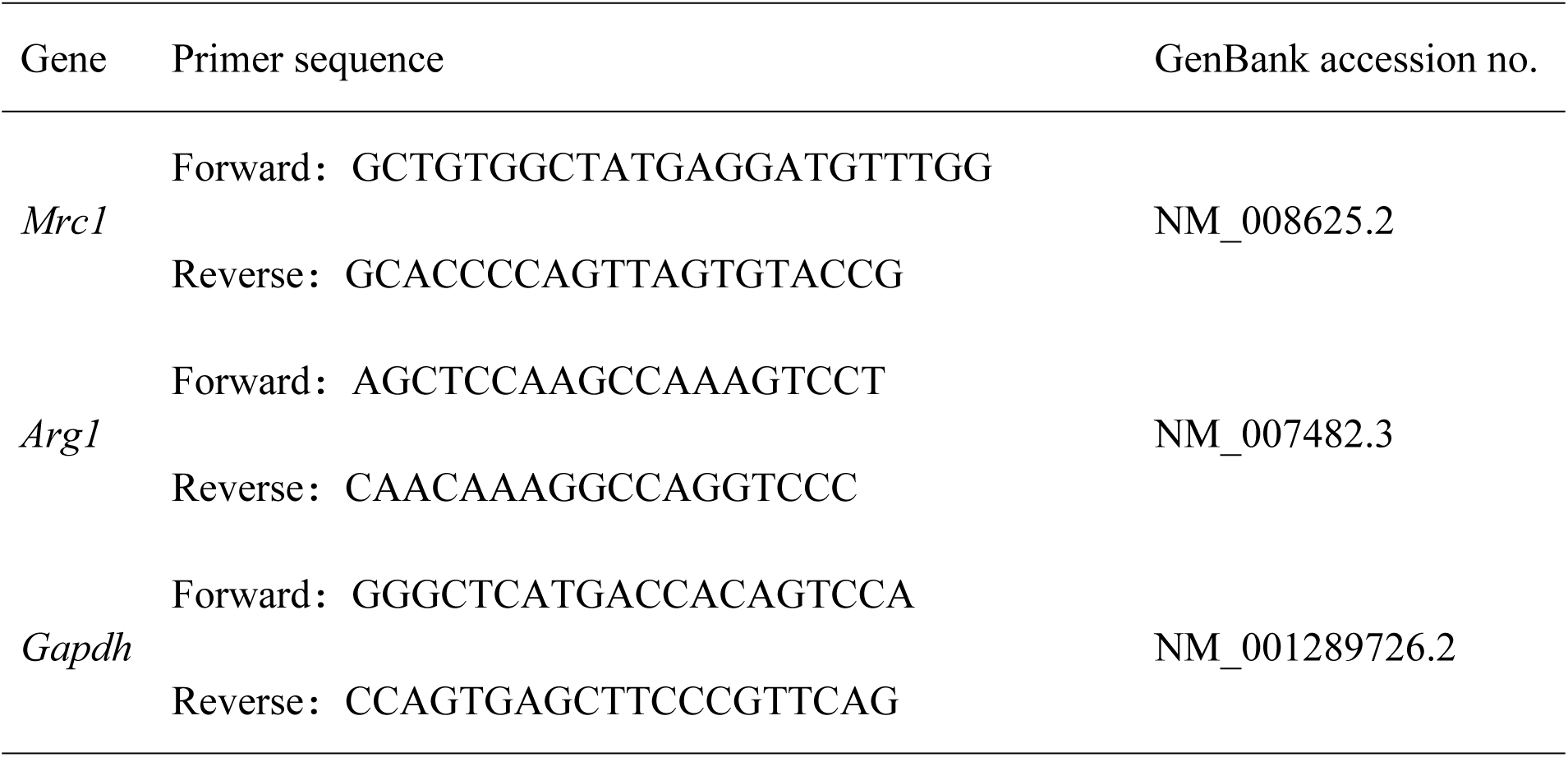
Primers for RT-qPCR.

### 2.12 Immunofluorescence (IF)

The expression of F4/80, CD86, and CD206 in lung tissue sections and cells were examined by IF. Briefly, the samples were permeabilized with 0.2% Triton X-100 and blocked with 5% BSA. The samples were then incubated overnight at 4 °C with primary antibodies anti-F4/80, anti-CD86, and anti-CD206 antibodies. After washing, the samples were incubated with secondary antibodies under light-protected conditions (20-25 °C), the cell nuclei were counterstained with DAPI for 10 minutes, followed by the addition of an anti-fade mounting medium. Finally, the samples were observed and imaged for analysis using fluorescence microscope.

### 2.13 Western Blot

After lysing lung tissue and cells, the total protein concentration was determined. Protein samples were subjected to 8/10% SDS-PAGE gel electrophoresis and then transferred to a PVDF membrane. The membrane was blocked with 5% skim milk and incubated with the primary antibody at 4°C for 10–12 h, CD206, Arg-1, JAK2, p-JAK2, STAT6, p-STAT6, PPAR-γ, and p-PPAR-γ. After the primary antibody incubation, the membrane was washed with TBST and then incubated with goat anti-rabbit or goat anti-mouse antibodies at room temperature with gentle agitation for 1 hour. Images were captured using an ECL Western blot detection system and subsequently analyzed using ImageJ software.

### 2.14 Molecular docking

The POG structures of small-molecule ligands were obtained from the PubChem database (http://pubchem.ncbi.nlm.nih.gov/) and converted into three-dimensional conformations using ChemOffice software, then saved in mol2 format. High-resolution crystal structures of AKT1, TNF, ESR1, STAT6, and BCL2 were downloaded from the RCSB PDB database (http://www.rcsb.org/), preprocessed using PyMOL software to remove water molecules and non-essential ligands, and saved as PDB files. Molecular docking was performed using AutoDock Vina 1.5.6. After preprocessing the ligand and protein—including hydrogenation, dehydration, and definition of torsional degrees of freedom—a docking grid was generated, and the optimal binding conformation was selected based on binding energy scores. Docking interactions were visualized and analyzed using Discovery Studio 2019 to generate a two-dimensional interaction diagram highlighting key residues and a three-dimensional ligand-binding model.

### 2.15 Statistical Analysis

All experimental data are presented as mean ± standard error (SEM). A one-way ANOVA was used to determine statistical differences between groups, with statistical significance set at P < 0.05. GraphPad Prism 9.0 was used for image quantification and statistical analysis.

## 3 Results

### 3.1 LC-MS-based Network Pharmacology analysis

LC-MS identified 62 constituents in SD/PM, primarily flavonoids, terpenoids, coumarins, and other compounds (**Fig. S1, Table 2**). A screening of the TCMSP database identified 32 constituents present in SD/PM, with 9 compounds overlapping with the LC-MS results (**Fig. 1A**).

**Fig. 1.**
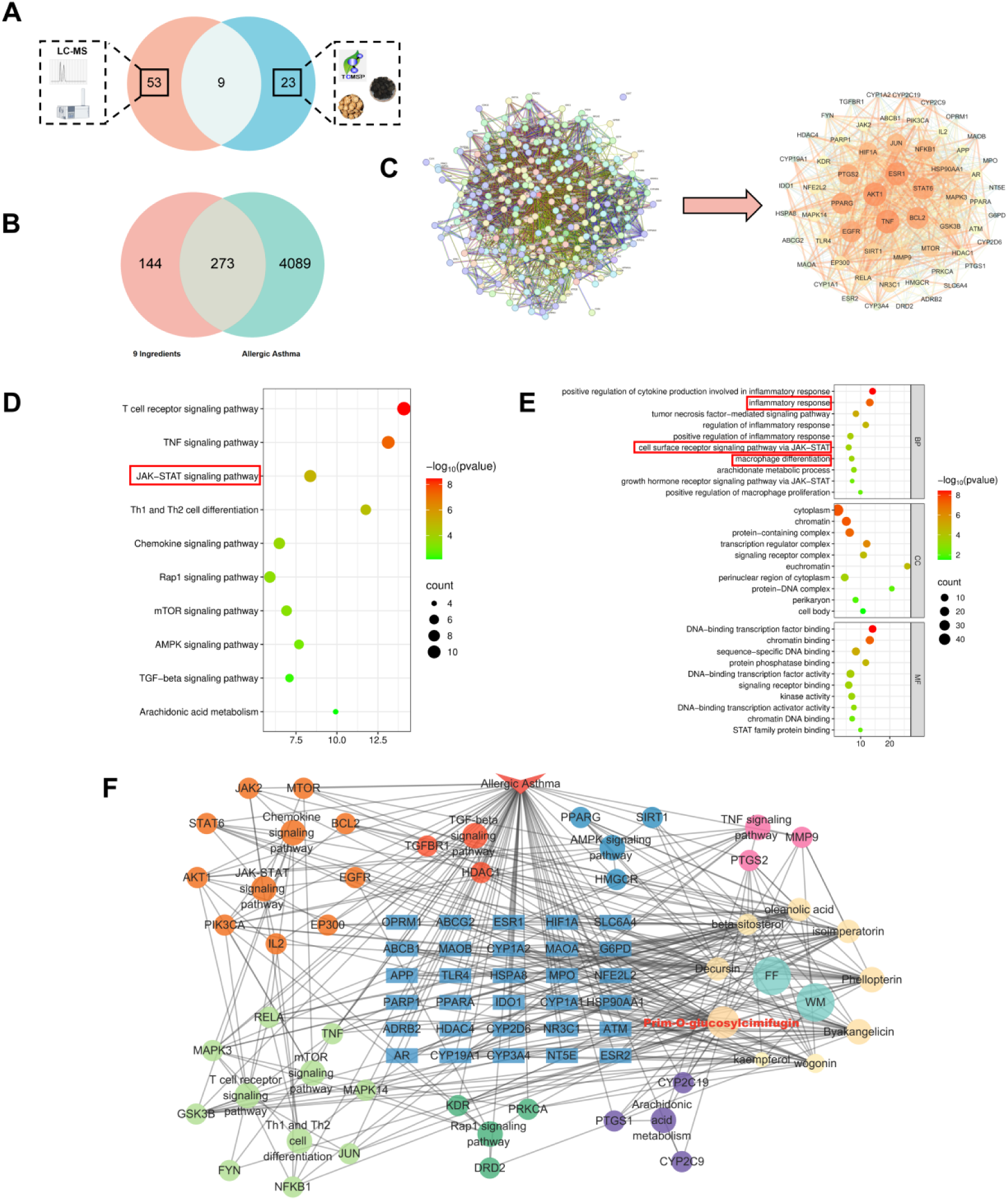
Network pharmacology analysis of SD/PM for the treatment of AA. (A) Venn diagram showing the intersection of LC-MS components and the database. (B) Venn diagram showing the intersection of drug components and disease targets. (C) PPI (protein-protein interaction) analysis. Each node represents a potential target for drug components to intervene in AA; their size and color are scaled from largest to smallest and from darkest to lightest, respectively, based on the node’s degree value. (D) GO functional enrichment analysis. (E) KEGG enrichment analysis. (F) “Drug-Compound-Disease-Pathway-Target” network diagram.

Databases related to disease and target prediction identified 4,362 AA-associated targets and 417 component-associated targets (**Fig. 1B**). Further analysis using a Venn diagram revealed 273 targets that were common to both the disease and component sets. A protein-protein interaction (PPI) network analysis was conducted using the STRING database in conjunction with the Cytoscape platform, identifying AKT1, TNF, ESR1, STAT6, and BCL2 as core targets (**Fig. 1C**). GO functional analysis revealed that the primary biological processes included inflammatory response, cell surface receptor signaling pathway via JAK-STAT, and macrophage differentiation; cellular components included cytoplasm, transcription regulator complex, and protein-DNA complex; and the primary molecular functions included DNA-binding transcription factor binding, protein phosphatase binding, and STAT family protein binding (**Fig. 1D**). KEGG analysis showed that pathways such as the T-cell receptor signaling pathway, TNF signaling pathway, JAK-STAT signaling pathway, and chemokine signaling pathway were significantly enriched (**Fig. 1E**). One study had confirmed that irisin drives M2 macrophage polarization by upregulating the JAK2-STAT6-PPARγ axis [10]. Using Cytoscape to construct a “drug–compound– disease–pathway–target” network, POG had the highest degree value (Degree = 27), followed by Phellopterin (Degree = 23), Byakangelicin (Degree = 23), oleanolic acid (Degree = 22), and other compounds (**Fig. 1F**).

Molecular docking analysis was performed between POG and the top five core targets. The lower the binding energy, the tighter the binding between the compound and the protein, and the higher the stability of the complex. The results show that POG exhibits good binding properties with key targets; the binding energies of POG with AKT1, TNF, ESR1, STAT6, and BCL2 are -9.1 kcal/mol, -6.1 kcal/mol, -5.3 kcal/mol, -6.0 kcal/mol, and -6.8 kcal/mol, respectively, all of which are below -5 kcal/mol (**Fig. 2A-E**). In summary, POG is the primary active component in SD/PM for alleviating AA, and its mechanism of action may be closely related to the immune-inflammatory response regulated by the JAK2-STAT6-PPARγ axis.

**Fig. 2.**
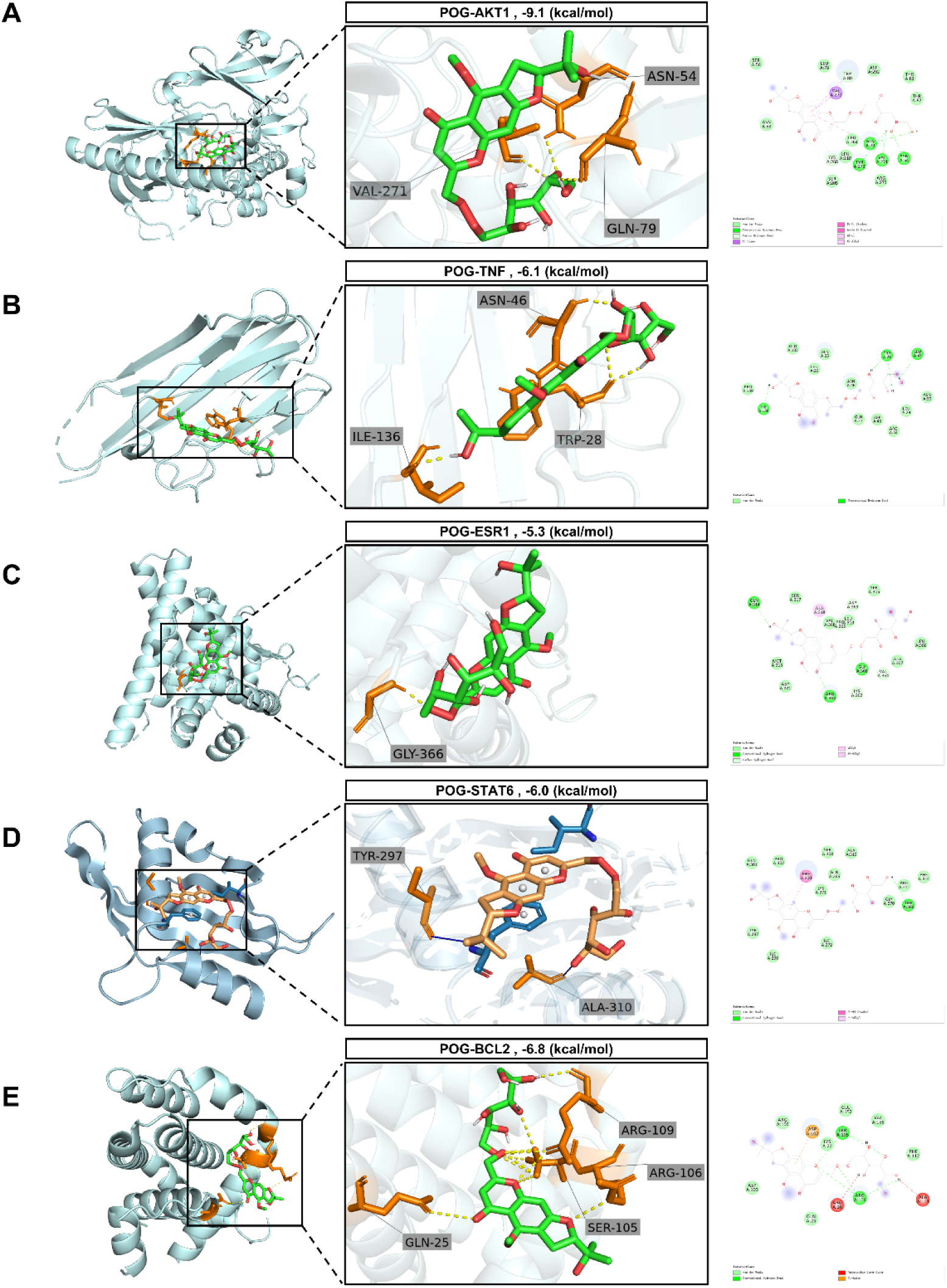
Molecular docking of POG and AA core targets. (A) Molecular docking of POG and AKT1. (B) Molecular docking of POG and TNF. (C) Molecular docking of POG and ESR1. (D) Molecular docking of POG and STAT6. (E) Molecular docking of POG and BCL2.

### 3.2 POG alleviates OVA-induced airway inflammation and lung injury in AA mice

OVA-induced AA mice were established to evaluate the effects of POG on AA (**Fig. 3A**). The results showed that the general condition of the mice in the model group was significantly abnormal compared to the control group, as evidenced by a significant decrease in body weight following OVA-induced modeling (**Fig. 3B**). At the same time, the lung index of the mice in the model group was significantly elevated, further indicating that their overall physiological status was markedly affected (**Fig. 3C**).

**Fig. 3.**
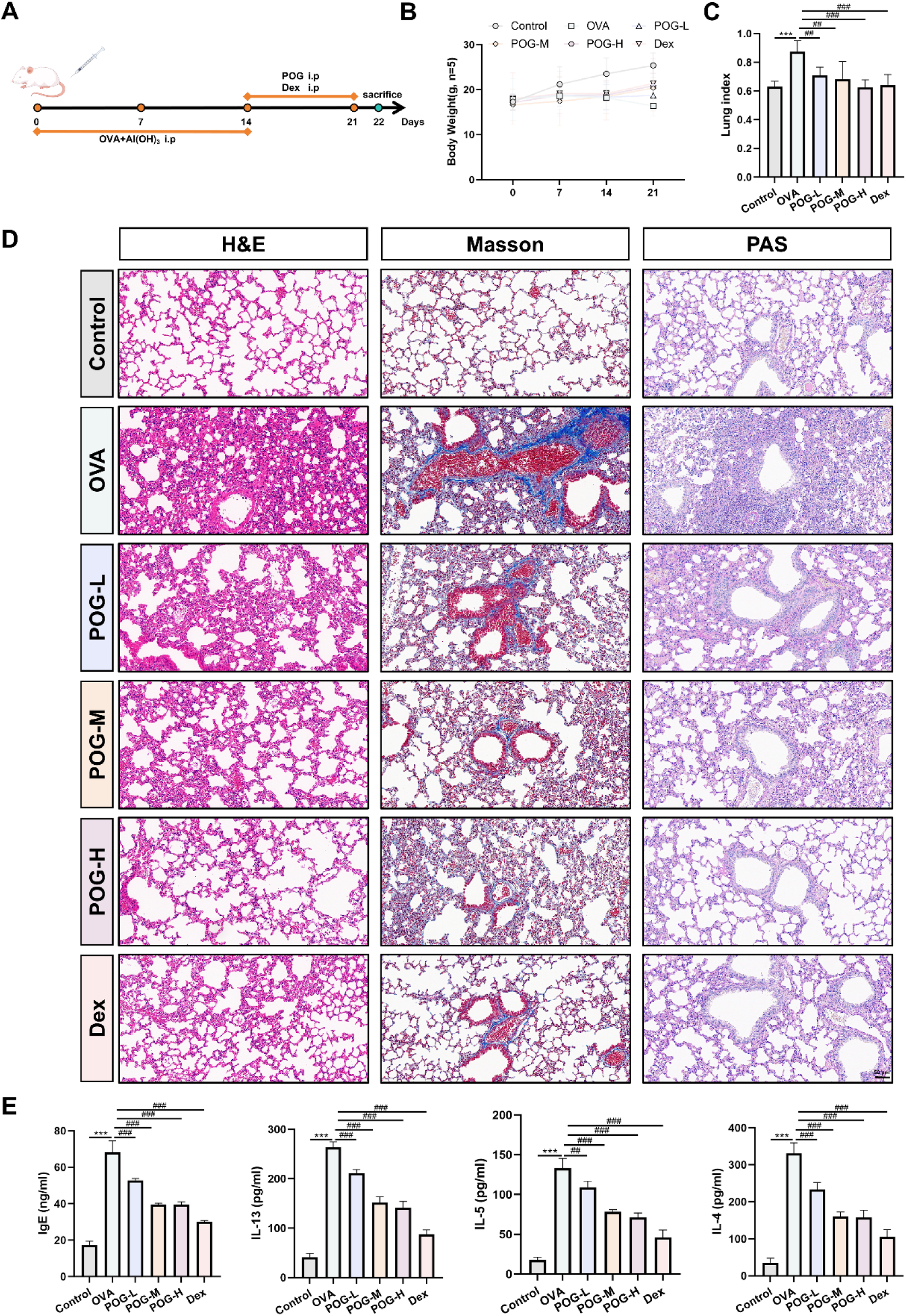
POG alleviates OVA-induced inflammatory damage in lung tissue. (A) Schematic diagram of the experimental design. (B) Changes in mouse body weight, n = 5. (C) Lung index in mice, n = 5. (D) H&E, Masson, and PAS histological staining of mouse lung tissue, n = 5. (E) Serum total IgE and type II cytokine levels in mice, n = 3. Compared with the Control group, ***p < 0.001; compared with the OVA group, ^###^p < 0.001, ^##^p < 0.01.

Studies have shown that AA is often accompanied by structural destruction and inflammatory damage to lung tissue [18]. Therefore, this study further analyzed pathological changes in lung tissue using histochemical staining (**Fig. 3D**). H&E staining revealed alveolar collapse, thickened alveolar walls, and inflammatory cell infiltration in the model group mice. Masson and PAS staining confirmed that, compared with the control group, collagen deposition in the lung tissue of the model group mice was significantly increased, suggesting airway remodeling. Treatment with POG markedly improved these pathological changes.

A growing body of evidence suggests that elevated levels of type II immune factors exacerbate AA [19–21]. To this end, this study used ELISA to measure total IgE and type II cytokine levels in mouse serum to evaluate the modulatory effect of POG on AA-related immune responses (**Fig. 3E**). The results showed that serum levels of IgE, IL-4, IL-5, and IL-13 in the model group mice were significantly higher than those in the control group. Intervention with POG and dexamethasone significantly reduced the levels of these inflammatory factors. In summary, POG can improve pulmonary tissue damage in AA by inhibiting Type II immune responses and reducing the release of inflammatory factors.

### 3.3 POG inhibits M2 macrophage polarization via the JAK2-STAT6-PPARγ pathway in AA mice

In this study, we analyzed macrophage subsets in lung tissue using flow cytometry to evaluate the regulatory role of POG in macrophage polarization. F4/80 was used to label total macrophages, while CD86 and CD206 were used to identify the M1 and M2 subtypes, respectively [22]. The results showed that the proportions of CD86⁺ and CD206⁺ macrophages were both significantly increased in the model group, suggesting that AA activated and enhanced the polarization of M1 and M2 macrophages (**Fig. 4A-B**). POG intervention significantly reduced the proportions of both M1 and M2 macrophages. Notably, POG’s inhibitory effect on the proportion of M2 macrophages was more pronounced than that on M1 macrophages, indicating that POG primarily alleviates the immune inflammatory response by regulating the polarization process of M2 macrophages. These findings were further validated at the genetic and protein levels using RT-qPCR, IF, and Western blot analysis. IF results demonstrated that POG significantly reduced the fluorescence expression levels of F4/80⁺CD86⁺ and F4/80⁺CD206⁺ in mice from the model group (**Fig. 4C-D**). RT-qPCR and Western blot results also showed that POG significantly reduced the gene and protein expression levels of the M2 macrophage markers CD206 and Arg-1 (**Fig. 4E**).

**Fig. 4.**
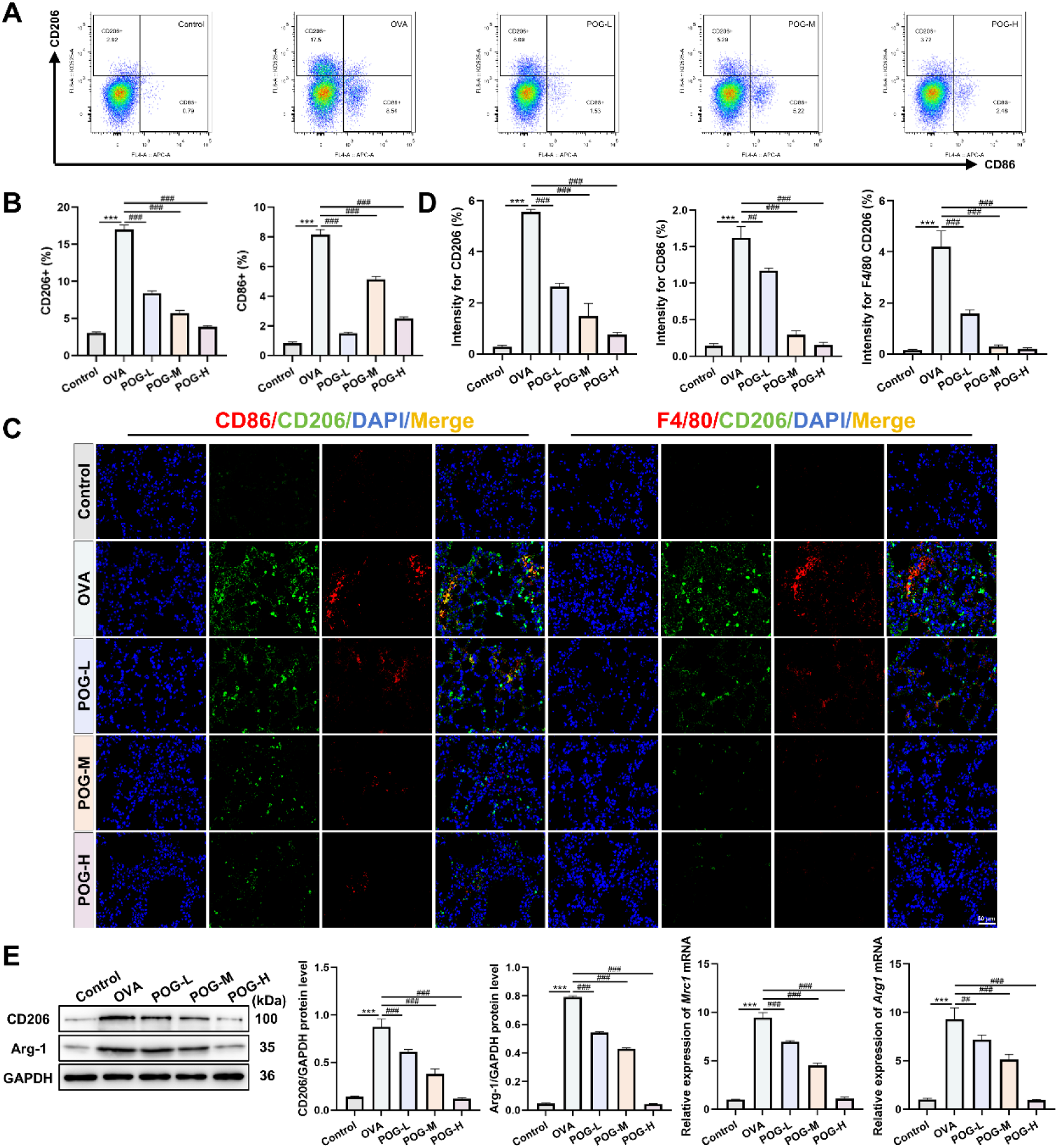
POG inhibits macrophage polarization. (A–B) Flow cytometric analysis of CD206^+^ and CD86^+^ cells in mouse lung tissue. (C–D) IF colocalization of CD86 with CD206 and CD206 with F4/80 in mouse lung tissue. (E) Western blot and RT-qPCR analysis of CD206 and Arg-1 levels in mouse lung tissue. Compared with the Control group, ***p < 0.001; compared with the OVA group, ^###^p < 0.001, n = 3.

Based on network pharmacology studies, the JAK2-STAT6-PPARγ axis is a key pathway through which POG improves AA. Therefore, we further used Western blot to detect the expression of key proteins in this signaling pathway. The results showed that, compared with the control group, the expression levels of p-JAK2/JAK2, p-STAT6/STAT6, and p-PPAR-γ/PPAR-γ were significantly elevated in the model group; following POG intervention, the expression levels of these proteins were significantly reduced (**Fig. 5A-B**). To further validate the regulatory role of this pathway in M2 macrophage polarization, this study employed the JAK2 agonist C-A1)for intervention. RT-qPCR and Western blot results showed that, compared with the POG treatment group, there was a trend toward upregulation of the CD206 and Arg-1 genes and proteins, indicating that C-A1 attenuated the effect of POG administration on M2 macrophage polarization (**Fig. 5C-D**). In summary, this study demonstrates that POG restores the immune-inflammatory balance in the lungs by inhibiting M2 macrophage polarization mediated by the JAK2-STAT6-PPARγ axis and partially inhibiting M1 polarization.

**Fig. 5.**
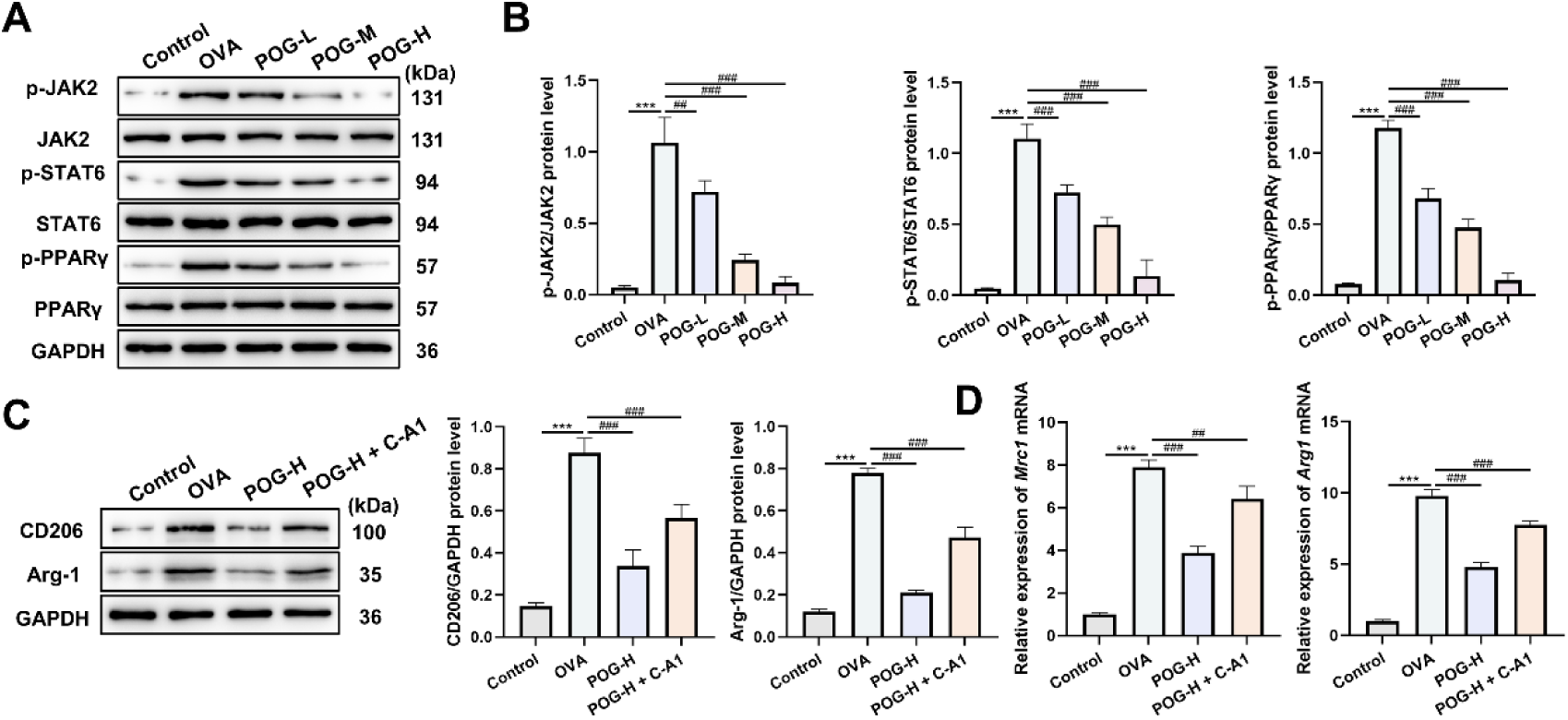
POG inhibits the JAK2-STAT6-PPARγ axis and reduces M2 macrophage polarization. (A–B) Protein expression levels of key factors in the JAK2-STAT6-PPARγ axis in mouse lung tissue. (C-D) Effects of the JAK2-STAT6-PPARγ pathway agonist C-A1 on the gene and protein levels of M2 polarization markers. Compared with the Control group, ***p < 0.001; compared with the OVA group, ^###^p < 0.001, ^##^p < 0.01, n = 3.

### 3.4 POG suppresses IL-4-induced M2 polarization in RAW264.7 cells by blocking JAK2-STAT6-PPARγ signaling

In this study, the cytotoxic effects of POG on RAW264.7 cells were first evaluated using the CCK-8 assay to determine the safe concentration range. The results showed that POG had no significant effect on cell viability within a concentration range of 0–8 μmol/L (**Fig. 6A**). Subsequently, an M2 polarization model was established in RAW264.7 cells induced by IL-4, and the cells were co-treated with different concentrations of POG (0, 2, 4, 8 μmol/L). The CCK-8 assay was used to further screen for the optimal concentration of POG for intervening in the inflammatory cell model. The results showed that (**Fig. 6B**), compared with the IL-4 model group, POG significantly restored cell viability within the concentration range of 2–8 μmol/L, with the most pronounced effect observed at 8 μmol/L. Flow cytometry results showed that, compared with the control group, the proportion of CD206⁺ macrophages significantly increased after IL-4 stimulation; following POG intervention, the proportion of CD206⁺ cells decreased in a concentration-dependent manner (**Fig. 6C**). Further RT-qPCR and Western blot results demonstrated that POG significantly reduced the mRNA and protein expression levels of CD206 and Arg-1 (**Fig. 6D-E**). IF results similarly demonstrated that POG significantly attenuated the IL-4-induced fluorescence intensity of CD206 (**Fig. 6F**). Furthermore, ELISA results showed that following POG treatment, levels of M2-associated chemokines CCL17, CCL22, and IL-10 were all significantly reduced (**Fig. 6G**).

**Fig. 6.**
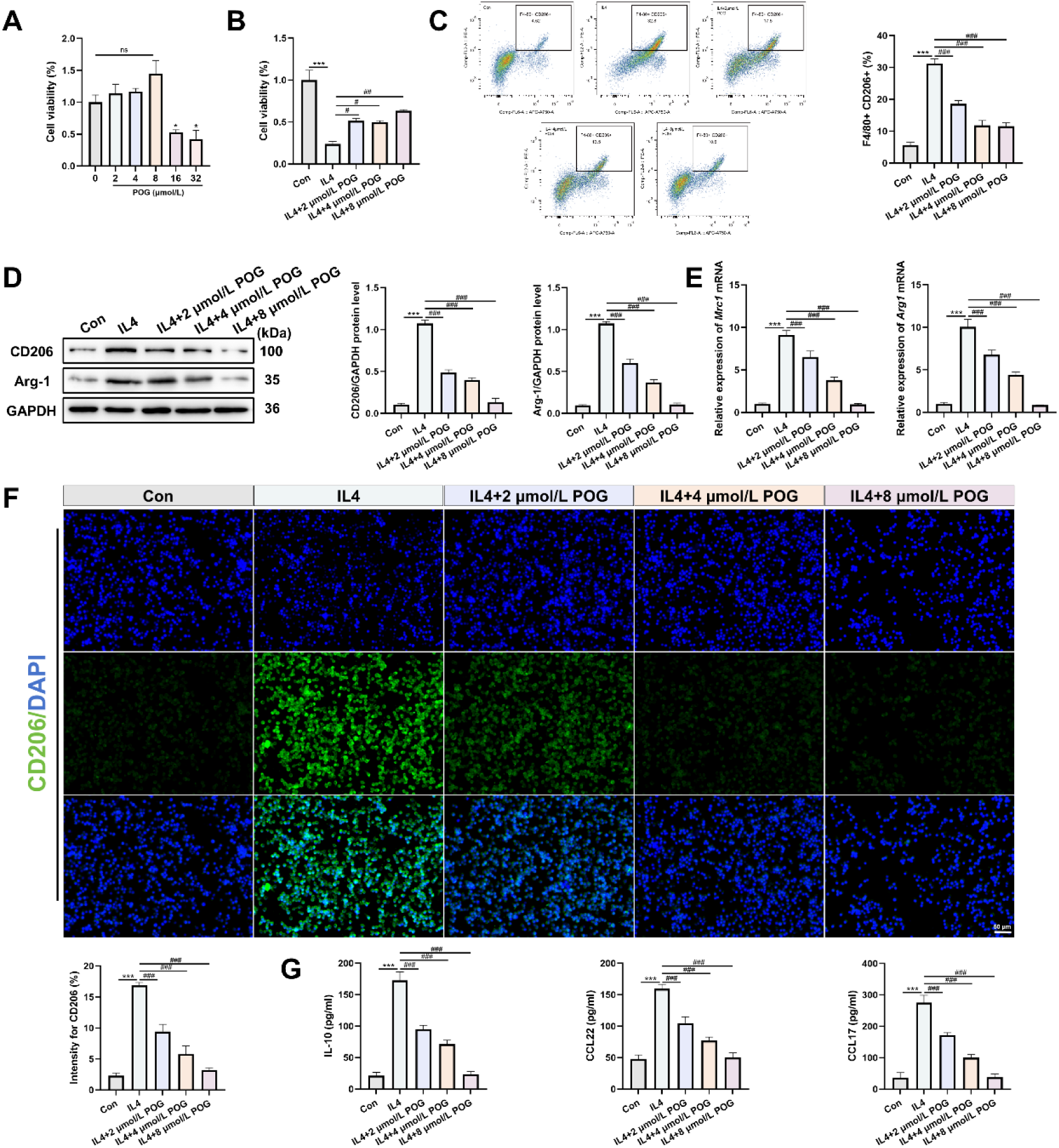
POG inhibits inflammatory responses and M2 polarization in RAW264.7 cells. (A-B) Effect of POG on RAW264.7 cell viability. (C) Flow cytometric analysis of CD206^+^ and F4/80^+^ cells in RAW264.7 cells. (D–E) Effect of POG on M2 polarization markers in RAW264.7 cells. (F) IF analysis of CD206 fluorescence intensity in RAW264.7 cells. (G) Effects of POG on M2-associated chemokines CCL17, CCL22, and IL-10 in RAW264.7 cells. Compared with the Con group, ***p < 0.001; compared with the IL-4 group, ^###^p < 0.001, ^##^p < 0.01, n = 3.

Additionally, this study utilized Western blot analysis to detect the levels of key proteins in the JAK2-STAT6-PPARγ axis, thereby validating the role of this pathway in vitro. The results showed that, compared with the control group, the protein expression levels of p-JAK2/JAK2, p-STAT6/STAT6, and p-PPAR-γ/PPAR-γ were significantly elevated after IL-4 stimulation; following POG intervention, the phosphorylation levels of these proteins decreased in a concentration-dependent manner (**Fig. 6H**). C-A1 exhibited a similar stimulatory effect. The results showed that, compared with the POG monotherapy group, both the mRNA and protein expression levels of CD206 and Arg-1 were significantly upregulated in the combination agonist group, indicating that C-A1 can partially reverse the inhibitory effect of POG on M2 polarization (**Fig. 6I-J**). In summary, this study demonstrates that POG exerts an immunomodulatory effect by inhibiting IL-4-induced M2 polarization of RAW264.7 macrophages through the blockade of the JAK2-STAT6-PPARγ signaling pathway (**Fig. 7**).

**Fig. 7.**
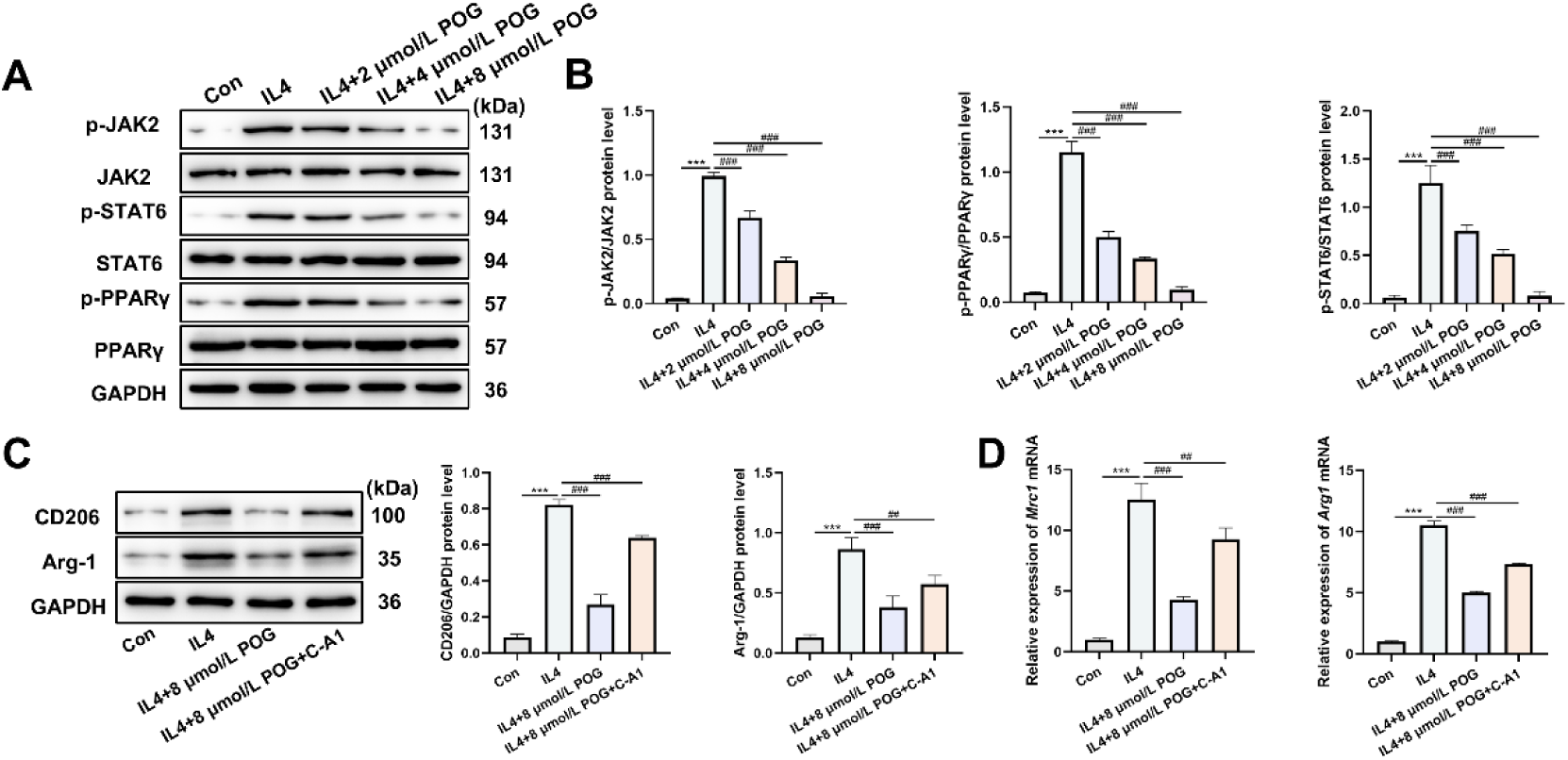
POG inhibits the JAK2-STAT6-PPARγ signaling pathway, thereby attenuating the M2 polarization process in RAW264.7 cells. (A-B) Protein expression levels of key factors in the JAK2-STAT6-PPARγ axis in RAW264.7 cells. (C-D) Effects of C-A1 on the gene and protein expression of M2 polarization markers in RAW264.7 cells. Compared with the Con group, ***p < 0.001; compared with the IL-4 group, ^###^p < 0.001, ^##^p < 0.01, n = 3.

**Fig. 8.**
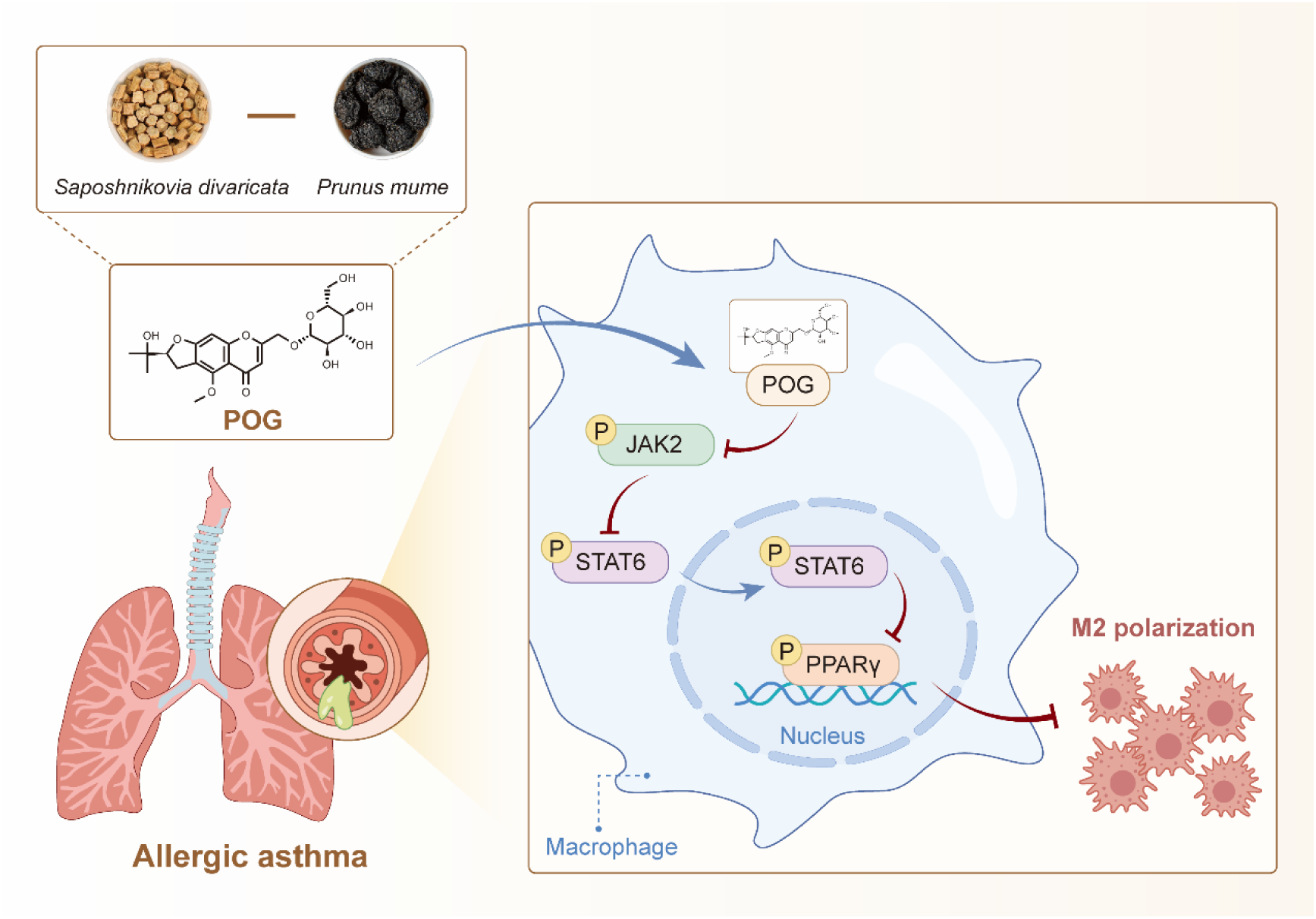
POG alleviates AA by inhibiting M2 polarization mediated by the JAK2-STAT6-PPARγ axis.

## 4 Discussion

AA is a severe chronic respiratory disease that poses a significant threat to human health[23]. Its core pathogenesis lies in the immune inflammatory response caused by the polarization of M2 macrophages [24]. Therefore, developing effective therapeutic interventions targeting the M2 macrophage polarization process is of great clinical significance for protecting lung function in patients with AA. Due to its multi-target, holistic regulatory properties, TCM has garnered widespread attention for its anti-inflammatory and immunomodulatory effects [25]. Based on this, this study utilized an OVA-induced AA mouse model and integrated network pharmacology, classical pharmacodynamics, and in vitro and in vivo molecular biology experiments to elucidate, from multiple perspectives, the therapeutic effects of the natural active compound POG on AA and its underlying molecular mechanisms.

Current network pharmacology research increasingly favors an approach that combines LC-MS identification with database searches [26]. Therefore, this study systematically investigated the active components and mechanisms of action of SD/PM in the treatment of AA by integrating LC-MS and network pharmacology techniques. Based on LC-MS detection results and matching analysis using the TCMSP database, a total of 9 major active components with anti-AA activity in SD/PM were identified. Among these, 6 components originated from SD and 3 from PM, suggesting that SD may be the primary contributor to the anti-AA effects of this formulation. PPI analysis revealed that AKT1, TNF, ESR1, STAT6, and BCL2 are key targets for the treatment of AA. Further KEGG pathway enrichment analysis indicated that the JAK-STAT signaling pathway was significantly enriched in the treatment of AA. Previous literature has reported that activation of the JAK-STAT6 pathway can induce ROS accumulation, lipid peroxidation, and ferrocytosis, thereby exacerbating asthma symptoms [27]. Network topology analysis identified POG as the key active ingredient in SD/PM responsible for alleviating AA. Notably, previous studies have confirmed that POG significantly alleviates LPS-induced acute lung injury in mice, further supporting its anti-inflammatory potential [28]. Molecular docking results further validated POG’s strong binding affinity for the aforementioned five core targets. In summary, POG may be the key active component in SD/PM for alleviating AA, and its mechanism of action may be closely related to the regulation of the JAK-STAT signaling pathway.

To systematically evaluate the therapeutic potential of POG for AA, this study conducted a comprehensive assessment of its in vivo efficacy through classical pharmacodynamic experiments. Typical pathological features of AA include alveolar damage, inflammatory cell infiltration in the airways, and airway wall remodeling, which exacerbate airway hyperresponsiveness and progressive decline in lung function in patients [29]. The histopathological results of this study showed that mice in the model group exhibited widespread destruction of alveolar structures, significant thickening of alveolar walls, and massive inflammatory cell infiltration, suggesting severe airway inflammatory responses and structural remodeling. Following POG intervention, all of the aforementioned histopathological lesions in the lung tissue were significantly alleviated. Studies have shown that M2 macrophage polarization promotes the release of type II cytokines such as IL-4, IL-5, and IL-13, which in turn drive B-cell class switching to produce IgE, synergistically exacerbating airway thickening and allergic inflammation [30,31]. Consistent with previous reports, the results of this study indicate that POG significantly reduces serum levels of IgE, IL-4, IL-5, and IL-13 in mice from the model group, suggesting that it may exert its anti-allergic effects by inhibiting the type II immune response associated with M2 macrophage polarization. Notably, the therapeutic efficacy of the POG-H group was comparable to that of dexamethasone group, indicating that POG has good potential for application in anti-asthma therapy and warrants further in-depth study. In summary, this study confirms that POG exhibits clear therapeutic efficacy in alleviating airway inflammation and alleviating lung tissue damage in AA.

Existing research indicates that the JAK-STAT signaling pathway plays a multifaceted role in the regulation of immune inflammation, particularly in the polarization of M2 macrophages [32]. Given the molecular complexity of this pathway, this study, building on previous findings, focuses on analyzing its key downstream effector molecules involved in the regulation of M2 macrophage polarization. One study has shown that M2 macrophage polarization depends on a synergistic signaling network formed by JAK2-STAT6 and PPAR-γ [10]. Based on this, we hypothesize that POG may exert its biological effects by interfering with this core signaling axis. In an OVA-induced AA mouse model, POG treatment significantly reduced the protein expression of p-JAK2, p-STAT6, and p-PPAR-γ in lung tissue, accompanied by a marked decrease in CD206 levels. The in vivo results were consistent with the in vitro validation experiments. Furthermore, this study employed the selective JAK2 inhibitor TQ05105 in intervention experiments to validate the aforementioned hypothesis. The results indicated that TQ05105 attenuated the effects of POG administration, consistent with the known regulatory role of this signaling axis in M2 macrophage polarization [33]. These findings suggest that POG has the potential to serve as an adjunct to glucocorticoids, allowing for a reduction in glucocorticoid dosage while maintaining the original therapeutic efficacy, thereby avoiding the systemic immunosuppression risks commonly associated with traditional glucocorticoid medications. In summary, this study confirms that POG exerts its beneficial effects on AA by inhibiting JAK2-STAT6-PPARγ-mediated M2 macrophage polarization.

However, this study still has some limitations. First, LC-MS can only reflect the relative abundance of chemical components in herbal medicines; it cannot fully reflect their absorption, metabolism, and contribution to pharmacological effects in the body. Second, validation of the mechanistic pathways still requires intervention studies targeting specific molecules through methods such as gene knockout or RNA interference. In addition, in the in vitro experiments of this study, only the RAW264.7 cell line was used. Further verification with primary or human macrophages is still required in the future.

## 5 Conclusion

In summary, the results of this study indicate that POG can effectively alleviate AA by reducing airway inflammation and lung tissue damage. Mechanistically, POG inhibits the polarization of M2 macrophages by downregulating the expression of key proteins in the JAK2-STAT6-PPARγ axis. This study provides new insights into the mechanism by which POG alleviates AA and offers solid evidence to further advance its clinical application.

## Ethics approval and informed consent

This study was approved by the Medical Ethics Committee of Haikou Traditional Chinese Medicine Hospital (Review Approval No.: **HHTCM2025-01-08**).

## CRediT

**Chong Li:** Conceptualization, Data curation, Funding acquisition, Writing – original draft – Preparation, Writing – review and editing – Preparation. **Hong Zhang:** Formal analysis, Investigation, Methodology. **Weiwei Yang:** Project administration, Resources. **Haiyan Fu:** Software, Supervision, Validation. **Honghong Huang:** Validation, Visualization.

## Clinical trial number

Clinical trial number: not applicable.

## Research methodologies involving AI tools

The authors report generative AI was not used in their research or preparation of this manuscript.

